# Core elements play distinct roles in promoter birth and transcriptional regulation

**DOI:** 10.64898/2025.12.02.691827

**Authors:** Syue-Ting Kuo, Wei-Yi Shen, Sheng-Wen Lai, Cheng-Wei Ni, Chih-Chiang Chang, Zoltan Palmai, Lee-Wei Yang, Nei-Li Chan, I-Ren Lee, Hsin-Hung David Chou

## Abstract

Gene expression shapes phenotypes and evolution. However, studies of gene regulation focus on transcription factors, overlooking core promoters. To investigate how promoters emerge and regulate transcription, we determined the sequence–function landscapes of core elements, −35 and −10, in constitutive and transcription factor-regulated promoters in *Escherichia coli*. Characterization of *in vivo* transcriptional landscapes and *in vitro* RNA polymerase–promoter interactions showed the −10 element as essential for promoter evolution from random sequences. In contrast, the −35 element, though broadly conserved, is dispensable for promoter birth. Instead, it exerts greater impact on gene regulation via coordinated interactions with transcription activators and RNA polymerase. We further showed that evolution fine-tunes the −35 and −10 sequences of transcription factor-regulated promoters to achieve near-maximal fold changes by lowering basal while elevating induced expression. A notable exception is P*_luxI_*, whose leaky expression provides a crucial baseline for initiating quorum sensing. These findings elucidate promoter design principles and underscore the interdependence and coevolution of core elements, RNA polymerase, and transcription factors.

## Introduction

Gene expression controls phenotypes, playing key roles in cell signaling, physiological acclimation, and evolution under changing environments ^1^. Promoters, comprising core elements recognized by RNA polymerase (RNAP) and accessory sites bound by transcription factors (TFs), provide the instruction to initiate gene expression. The interactions among promoters, RNAP, and TFs collectively determine the timing, location, and magnitude of RNA synthesis. Understanding how promoter sequence variation affects transcriptional output is essential for dissecting gene regulatory mechanisms ^2^, engineering genetic circuits for synthetic biology ^3^, uncovering the genetic basis of speciation and developmental innovation ^4,5^, and fundamentally, deciphering how new promoters emerge through sequence evolution ^6–9^.

Bacterial promoters consist of multiple core elements—UP, −35, spacer, extended −10, −10, discriminator, core recognition, and start—arranged in the 5′-to-3′ order ^2,10,11^. Transcription begins with a σ factor, primarily the housekeeping σ^70^, that assists RNA polymerase (RNAP) in recognizing the −35 and −10 elements ^12^. Subsequently, RNAP unwinds DNA from the −10 element and initiates RNA synthesis at the downstream transcription start site (Fig. 1a) ^13^. Among core promoter elements, prior sequence and functional analyses indicate that the −35 and −10 elements are the most conserved and critical for transcription initiation, with TTGACA and TATAAT representing their respective consensus sequences ^11,14–17^. The evolutionary significance of the −35 and −10 elements is further underscored by prior studies where random or genomic sequences were synthesized for assessing and evolving promoter activity ^7–9^. Among these, Yona et al. observed the recurrent emergence of the −35 and −10 motifs (TTGnnn and TAnnnT, respectively), but not of other core elements, in *de novo* promoters evolved from random sequences ^7^. Notably, some *de novo* promoters contained only the −10 element without a recognizable −35 motif. This finding supports the hypothesis that the −35 element may be nonessential to promoter birth and transcription initiation, given its higher variability among native promoters and that its contribution to promoter activity can be replaced by the extended −10 element ^18–20^. However, it challenges the current structural model in which −35 sequence recognition by the σ^70^ domain 4 (σ^70^_4_) is critical for initial RNA polymerase–promoter binding ^13,21–23^.

**Figure 1.**
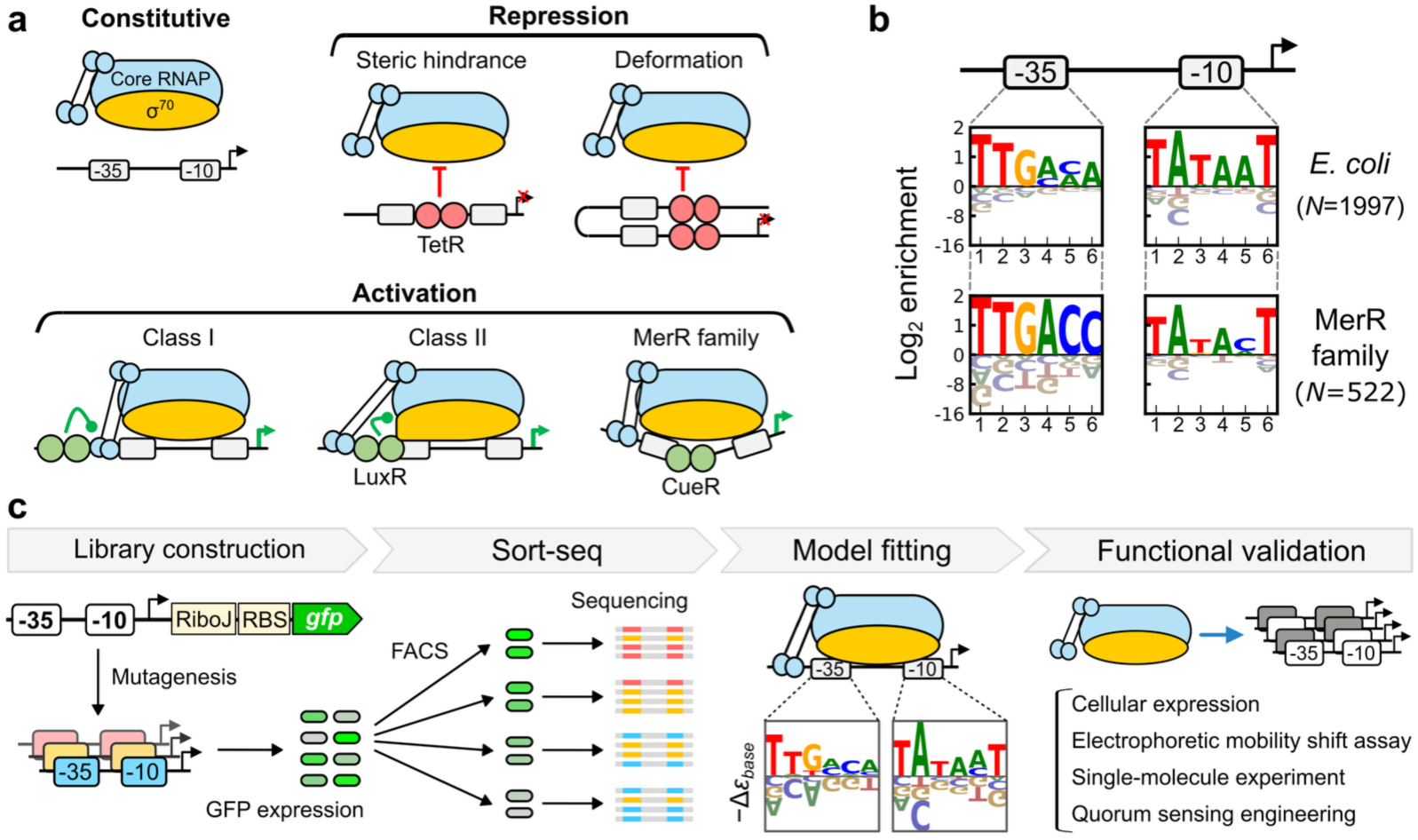
Dissecting the roles of the −35 and −10 elements in promoter birth and regulation. **a** Promoter types—depicting RNA polymerase (RNAP), transcription factors (TFs), and promoter interactions. TFs characterized in this study (TetR, LuxR, CueR) are indicated, with repressors and activators represented by red and green circles, respectively. **b** Sequence enrichment of the −35 and −10 elements in *E. coli* σ^70^-dependent promoters and promoters regulated by the MerR family across diverse species ^11,28^. **c** Research workflow, including construction of promoter libraries, sort-seq quantification of promoter strength, model fitting to estimate the free energy contribution of each nucleobase (*−Δε_base_*) in the −35 and −10 elements, and functional validation of promoter variants. RiboJ ^82^, self-cleaving ribozyme to make mRNA with a uniform 5′ end; RBS, ribosome binding site; *gfp*, green fluorescent protein gene; FACS, fluorescence-activated cell sorting.

Interestingly, whereas constitutive promoters generally display high −35 and −10 sequence conservation ^11,14,24,25^, promoters regulated by certain TFs exhibit distinct conservation patterns at these elements, suggesting that TF-mediated regulation may also depend on core promoter properties (Fig. 1b) ^2,26–28^. In terms of their influences on gene expression, bacterial TFs can be broadly classified as repressors or activators (Fig. 1a). Repressor TFs typically bind within the 60 bp upstream and downstream of the transcription start site ^29^. They prevent RNAP from generating transcripts by directly occupying the core promoter (steric hindrance, e.g., TetR), looping DNA for promoter seclusion (deformation, e.g., LacI), or binding downstream to impede transcriptional elongation (roadblock, e.g., TetR) ^2,30^. Repression is most effective when their binding sites (TFBSs) overlap with the spacer element ^31,32^. Engineering of repressor-regulated promoters further identified two attributes for achieving high induction folds: (1) high-affinity TFBSs to reduce leaky expression ^33,34^; and (2) near-consensus −35 and −10 elements to confer strong expression upon derepression ^33,35^.

In contrast to repressors, activator TFs involve specific interactions with RNAP, reflected by the spatial restriction of their TFBSs (Fig. 1a): within the UP element (Class I; e.g., Crp), partially overlapping the UP and −35 elements (Class II; e.g., LuxR), or partially overlapping the −35 and spacer elements (MerR family; e.g., CueR) ^27^. Class I activation involves direct contact between activator TFs and the C-terminal domain of the RNAP α subunit (αCTD)^36^, whereas Class II activation primarily depends on interactions with σ^70^ ^37,38^. Prior studies have reported reduced - 35 sequence conservation among Class II promoters, interpreted as an adaptation to minimize basal transcriptional activity ^27,38,39^. By contrast, MerR-type promoters feature a near-consensus −35 element and a less conserved −10 element (Fig. 1b), separated by an overlength spacer element (19-20 bp; typical range: 16-18 bp) that disfavors RNAP binding ^40,41^. MerR-type TFs activate transcription through a unique mechanism ^42,43^: they bind to TFBSs in both basal and induced states, and upon allosteric induction by cognate ligands, ranging from metal ions to antibiotics and redox-active small molecules, bend the DNA to restore proper spacing and spatial orientation between the −35 and −10 elements ^44,45^. The extended spacer length in MerR-type promoters is evidently an adaptation to this mode of action ^41^. However, the functional significance of their contrasting −35 and −10 sequence conservation remains poorly understood.

These observations from both activator- and repressor-regulated promoters implicate core promoter elements as integral components of transcriptional regulation. However, except for limited instances ^33,35,46–48^, prior research primarily focuses on TFs, TFBSs, inducer molecules, and their interactions ^49–59^, with much less attention given to how variation in core promoter elements modulates transcription. To unravel the design principles of bacterial promoters and transcriptional regulation, we performed sort-seq experiments to systematically determine the sequence–function relationships of the −35 and −10 elements in two constitutive promoters and three TF-regulated promoters in *Escherichia coli*, using cellular expression of the green fluorescent protein (GFP) as a proxy for promoter strength (Fig. 1c). We then applied biophysical modeling, performed bioinformatics analyses, and conducted physiological and biochemical experiments for data analysis and functional validation. This integrative framework allows us to quantitatively distinguish the contributions of the −35 and −10 elements to promoter birth and transcriptional regulation from genomic, biochemical, and evolutionary perspectives.

## Results

### The −10 element predominantly shapes the functional landscape of constitutive promoters

Expression of constitutive promoter libraries PL_C17_ and PL_C16_, bearing fixed 17- and 16-bp spacers and all possible −35 and −10 sequences (4^12^ =16,777,216), was measured in two sort-seq replicates (Fig. 2a and Supplementary Fig. S1a). Throughout this study, promoter strength is expressed in relative fluorescence units (RFU), which correlate linearly with mRNA levels as shown previously and confirmed here (Fig. 4d) ^11^. PL_C17_ was synthesized and characterized as a single pool previously ^11^, whereas PL_C16_ was constructed and examined as four complementary sub-libraries (4^11^ each) to increase the sampling depth in this work. Two sort-seq replicates collectively detected 11,670,734 (69.6%) variants in PL_C17_ and 14,403,094 (85.8%) variants in PL_C16_, with 7 and 16 median reads per variant, respectively (Supplementary Fig. S2a). We retained 1,022,018 PL_C17_ variants that passed a data quality filter (≥ 20 reads and detection in at least two sort-seq ranks) for library property analysis (Pearson’s *r* = 0.90 between replicate measurements; Fig. 2b). By contrast, despite the higher sampling depth of PL_C16_, its left-shifted promoter strength distribution (1.2–247 RFU, compared with 2.2–1,115 RFU for PL_C17_) placed more variants below the detection limit (6.0 RFU, set by cell autofluorescence; Fig. 2c). The marked difference in promoter strength distributions underscores the distinct promoter contexts of PL_C17_ and PL_C16_, strengthening the representativeness of our experimental design. To ensure data quality, we applied an additional filter to retain PL_C16_ variants with <5-fold difference between replicate measurements, resulting in 213,081 variants with moderate correlation (Pearson’s *r* = 0.64; Fig. 2b).

**Figure 2.**
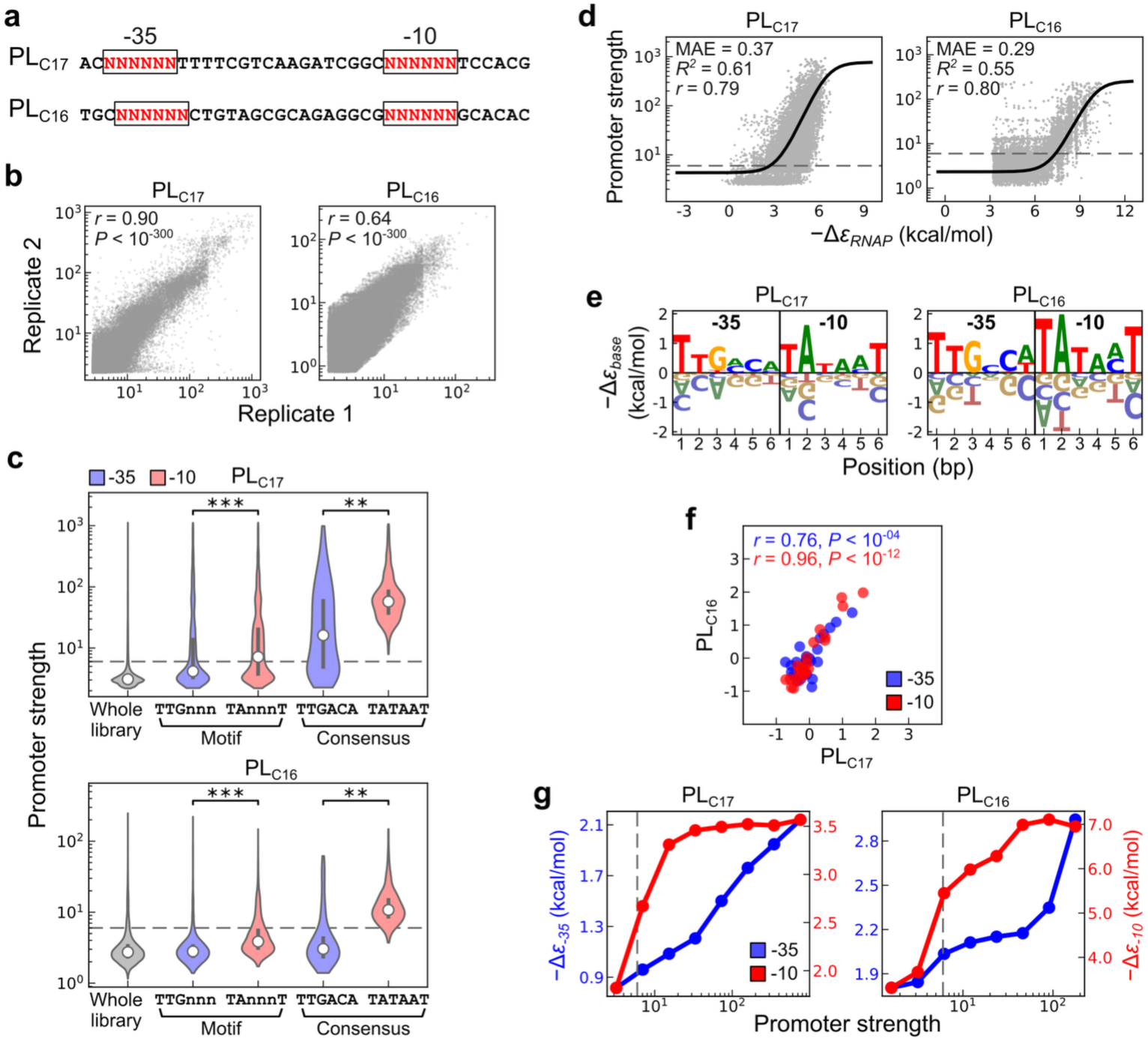
Functional contribution of the −35 and −10 elements in constitutive promoters. **a** Sequence composition of constitutive promoter libraries. Randomized base positions are shown in red. **b** Correlation (Pearson’s *r*, *t*-test) between promoter strength measured in two sort-seq replicates. Promoter strength is reported as the relative fluorescence unit. **c** Promoter strength distributions of the whole library and of variants bearing the −35 and −10 motifs or consensus sequences. Medians and interquartile ranges are indicated by white circles and vertical lines, respectively. Statistical significance from the Mann–Whitney U test is indicated as follows: \*\*\**P* < 10^−100^; \*\**P* < 10^−10^. **d** Model fitting performance. Curves and dots represent model predictions and experimental data, respectively. *−Δε_RNAP_*, RANP–promoter binding energy; MAE, mean absolute error. **e** Model-estimated free energy contribution of each nucleobase (*−Δε_base_*) in the −35 and −10 elements. **f** Correlation (Pearson’s *r*, *t*-test) of *−Δε_base_* between PL_C17_ and PL_C16_ (*N*_-35_ = 24, *N*_-10_ = 24). **g** Relationship between promoter strength and the binding energy contributed by the −35 (*−Δε_-35_*) and −10 (*−Δε_-10_*) elements. Variants in PL_C17_ and PL_C16_ are grouped into eight equal-sized bins across the whole-library promoter strength distributions shown in **c**. Dots indicate the group means of *−Δε_-35_* and *−Δε_-10_*. In **c**, **d**, and **g**, dashed lines indicate the promoter strength detection limit owing to cell autofluorescence.

**Figure 3.**
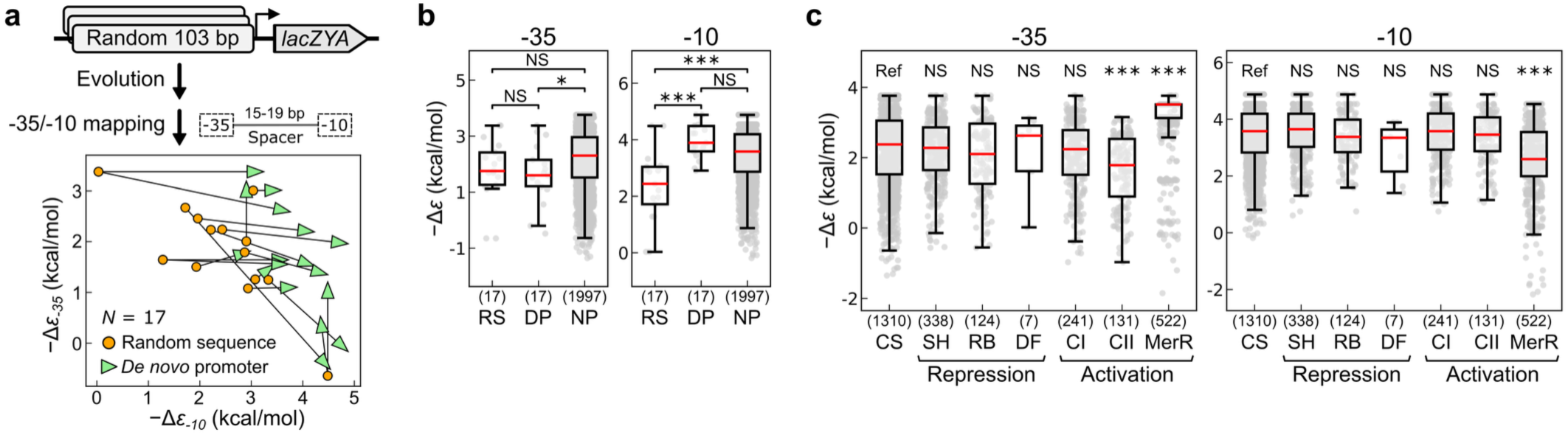
Functional contributions of the −35 and −10 elements in native and *de novo* promoters. **a** Evolutionary shifts in the binding energy contributions of the −35 (*−Δε_-35_*) and −10 (*−Δε_-10_*) elements. A prior study evolved random sequences to drive expression of the *lac* operon ^7^. The −35 and −10 elements in these sequences are predicted by the PAS model. Lines represent mutation paths that connect random sequences to their derived *de novo* promoters. **b** Distributions of *−Δε_-35_* and *−Δε_-10_* among random sequences (RS) and *de novo* promoters (DP), compared with *E. coli* native promoters (NP) annotated in RegulonDB ^68^. **c** Distributions of *−Δε_-35_* and *−Δε_-10_* among constitutive (CS), repressor-regulated (SH: steric hindrance, RB: roadblock, DF: deformation), and activator-regulated (CI: Class I, CII: Class II) promoters in *E. coli* ^68^ and in promoters regulated by the MerR family across diverse species ^28,83^. Constitutive promoters serve as the reference (Ref) for statistical comparisons. In **b** and **c**, dots, boxes, whiskers, and red bars represent individual promoters, interquartile ranges, 1.5-fold interquartile ranges, and medians, respectively. Sample sizes are shown in parentheses. Statistical significance from the Mann–Whitney U test is indicated as follows: \*\*\**P* < 0.001; \**P* < 0.05; NS, not significant.

**Figure 4.**
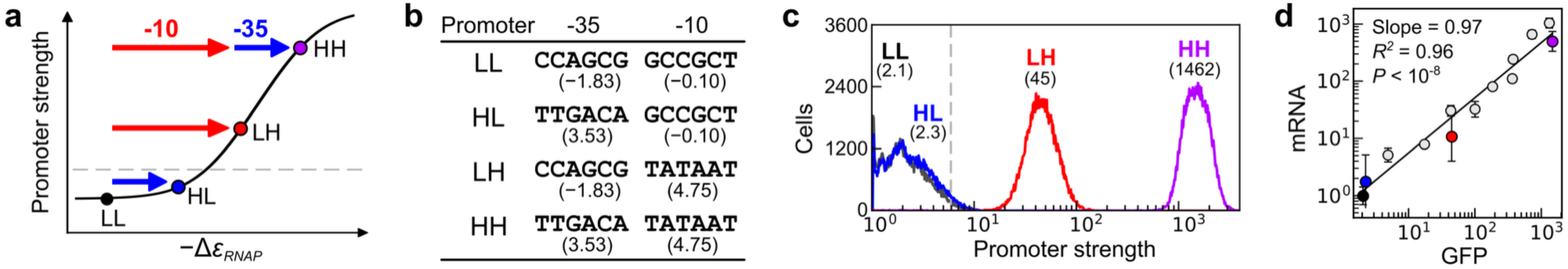
*In vivo* validation of the essentiality of the −35 and −10 elements for promoter activity. **a** Conceptual model showing the nonlinear correspondence between *−Δε_RNAP_* and promoter strength. **b** Promoter variants comprise combinations of −35 and −10 sequences differing in binding energy (kcal mol^-1^; shown in parentheses) and a transcription-promoting spacer (5′-TTTGCTTTAAATGGGG-3′) ^11^. **c** Promoter strength of the four variants quantified by flow cytometry (50,000 cells each). **d** Linear regression of mRNA fold-changes against GFP levels (*i.e.*, promoter strength) for the four promoter variants (colored as in **c**) and the spike-in variants (white circles; Supplementary Table S3). The lowest-expressing variant (LL) serves as the reference for calculating mRNA fold changes. Dots and error bars represent means and standard deviations from at least three independent measurements. In **a** and **c**, dashed lines indicate the promoter strength detection limit (*i.e.*, 6.0 RFU).

We assessed the relative significances of the −35 and −10 elements in constitutive promoters using motif- and sequence-level analyses. In the motif-level analysis, we compared the promoter strength of four subsets from PL_C17_ and PL_C16_: those carrying the canonical −35 (TTGnnn) or −10 (TAnnnT) motifs, and those carrying the −35 or −10 consensus sequences (Fig. 2c). Relative to the whole library, the promoter strength distributions of these subsets were all significantly higher in both PL_C17_ and PL_C16_ (*P* < 10^−10^, Mann–Whitney U test). Nevertheless, the magnitudes differed—subsets with the −10 motif generally showed higher promoter strength than those with the −35 motif (*P* < 10^−100^), and the −10 consensus overall exceeded the −35 consensus (*P* < 10^−10^). The greater importance of the −10 element over the −35 element in defining promoter identity was evident: in PL_C17_ and PL_C16_, 100% and 96% of variants with the −10 consensus exhibited detectable transcription activity, compared with only 71% and 20% of variants with the −35 consensus.

In the sequence-level analysis, we applied “Promoter Architecture Scanner (PAS) ^11^,” a thermodynamic model that infers the quantitative relationship between promoter sequence and strength by estimating the per-base free energy contributions (*−Δε_base_*) of the −35 and −10 elements to RNAP–promoter binding (Supplementary Fig. S3). Throughout, free energies (normally negative) are sign-inverted for intuitive interpretation, such that higher values indicate stronger binding. To minimize sampling bias from the right-skewed promoter strength distribution, we divided each library into ten equal-sized bins and randomly sampled 2,000 variants per bin for model fitting ^60^. The resulting PL_C17_ and PL_C16_ models yielded performance metrics of *R*^2^ = 0.61 and 0.55, Pearson’s *r* = 0.79 and 0.80, and mean absolute error (MAE) = 0.37 and 0.29, respectively (Fig. 2d). Model-estimated *−Δε_base_* correlated significantly (*P* < 10^-4^) between the two libraries, with Pearson’s *r* = 0.76 and 0.96 for the −35 and −10 elements, respectively (Fig. 2e, f). We compared the contributions of the −35 and −10 elements to promoter strength by dividing PL_C17_ and PL_C16_ variants into eight promoter strength groups and computing the group mean of *−Δε_-35_* and *−Δε_-10_*, defined as the *−Δε_base_* sum of each element (Fig. 2g). *−Δε_-10_* rose sharply from undetectable to low-expression levels in both libraries, whereas *−Δε_-35_* increased gradually with promoter strength in PL_C17_ and more sharply in PL_C16_ from moderate- to high-expression levels. Together, results from two constitutive promoter libraries suggest that the dominant effect of the −10 element makes it essential to promoter evolution from transcription-inactive sequences.

### Quantitative analysis shows promoter birth requires the −10 rather than the −35 element

To test the essentiality of the −10 element for promoter identity and evolution, we applied the PAS model to predict the −35 and −10 elements and estimate their free energy contributions in *de novo* promoters evolved from random sequences. The PAS model is well-suited for this task, as it has been shown to reliably map the two promoter elements across diverse bacteria ^11^. We analyzed 17 *de novo* promoters evolved from 14 103-bp random sequences in an evolution experiment ^7^, in which *E. coli* could grow only if these sequences drove the *lac* operon expression (Fig. 3a). The PAS model, trained on the PL_C17_ dataset, scanned both random sequences and *de novo* promoters to identify the core promoter region, where the −35 sequence, −10 sequence, and spacer length jointly maximized the RNAP–promoter binding energy (*−Δε_RNAP_*; see Methods). Comparison of random sequences with their derived *de novo* promoters showed that promoter birth was driven predominantly by significant increases in *−Δε_-10_* rather than *−Δε_-35_* (*P* < 0.001; Fig. 3a, b). Notably, through just one mutation step, *−Δε_-10_* among *de novo* promoters became indistinguishable from that of *E. coli* native promoters, supporting the idea that the −10 element functions as an “ON/OFF switch” for promoter identity (Fig. 3b). In contrast, *−Δε_-35_* among *de novo* promoters was indistinguishable from random sequences and remained lower than in native promoters (*P* < 0.05; note that three mutations in Fig. 3a increased *−Δε_-35_*). This suggested that the −35 element was dispensable for promoter birth but might act as a “volume tuner” to adjust promoter strength in subsequent evolution. Notably, changes in *−Δε_-35_* and *−Δε_-10_* during the evolution experiment mirrored those across the PL_C17_ and PL_C16_ sequence–function landscapes (Fig. 2g). Collectively, these results support an audio system–like design of bacterial promoters, which prompted us to investigate its quantitative and biochemical basis, as well as its implications for transcriptional regulation.

### Audio system–like behavior results from −10 dominance and nonlinearity between *−Δε_RNAP_* and transcription

Based on the above observations, we proposed that the audio system–like behavior of bacterial promoters emerged from (1) the greater functional contribution of the −10 relative to the −35 element and (2) the sigmoidal relationship between *−Δε_RNAP_* and promoter strength determined by the thermodynamic principle (Fig. 2d) ^15,35,61^. Owing to these, a −10 element created by random mutations should be essential and may suffice for promoter identity, whereas a *de novo* −35 element alone is likely insufficient to raise *−Δε_RNAP_* above the threshold for transcription initiation (Fig. 4a). We tested this hypothesis *in vivo* by constructing four promoter variants (LL, HL, LH, HH), representing the combinations of two −35 and two −10 sequence variants, each with low (L) or high (H) free energy contributions (Fig. 4b). The test was performed in two promoter contexts: one with a 16-bp spacer sequence lacking the extended −10 element (Fig. 4b) ^11^, and the other with a 17-bp spacer containing the extended −10 element (Supplementary Fig. S4a) ^62^, both of which promoted transcription and were identified in prior studies. The purpose of using transcription-promoting spacers here was to assess the essentiality, rather than the sufficiency, of the −35 or −10 elements for promoter identity. In both contexts, LH promoters with low *−Δε_-35_* and high *−Δε_-10_* (i.e., −10 consensus) showed detectable expression, though lower than HH promoters (Fig. 4c and Supplementary Fig. S4b). In contrast, HL promoters with high *−Δε_-35_* (i.e., −35 consensus) and low *−Δε_-10_* exhibited expression levels comparable to LL promoters. Confirmed at both mRNA and GFP levels (Fig. 4d and Supplementary Fig. S4c), the *in vivo* results recapitulate the switch- and tuner-like roles of the −10 and −35 elements in controlling promoter strength, respectively, supporting the “-10 essential and −35 dispensable” hypothesis.

To specifically probe RNAP interactions with the −35 and −10 elements, we selected the LL, HL, LH, and HH promoters lacking the extended −10 element for *in vitro* characterization, because its functional compensation for the −35 element, demonstrated previously ^18–20^, could confound the interpretation. Using the electrophoretic mobility shift assay (EMSA) to examine whether RNAP binding to the four promoters (each at 0.25 µM each) can transition into the productive open complex (RPo), we observed three distinct shifted bands in addition to free DNA, likely corresponding to the RPo state and two non-productive binding forms (Fig. 5a and Supplementary Fig. S5). The strongly expressed HH promoter predominantly yielded RPo at RNAP concentrations above 0.25 µM. In contrast, the transcription-silent LL and HL promoters mainly produced the two non-productive complexes, represented by a faster-migrating band at 0.25 µM RNAP and a slower-migrating band at 1.0 µM RNAP. Consistent with the *in vivo* results (Fig. 4c), only the transcription-active HH and LH promoters produced an appreciable amount of RPo, with the weaker LH promoter producing a larger fraction of the two non-productive complexes.

**Figure 5.**
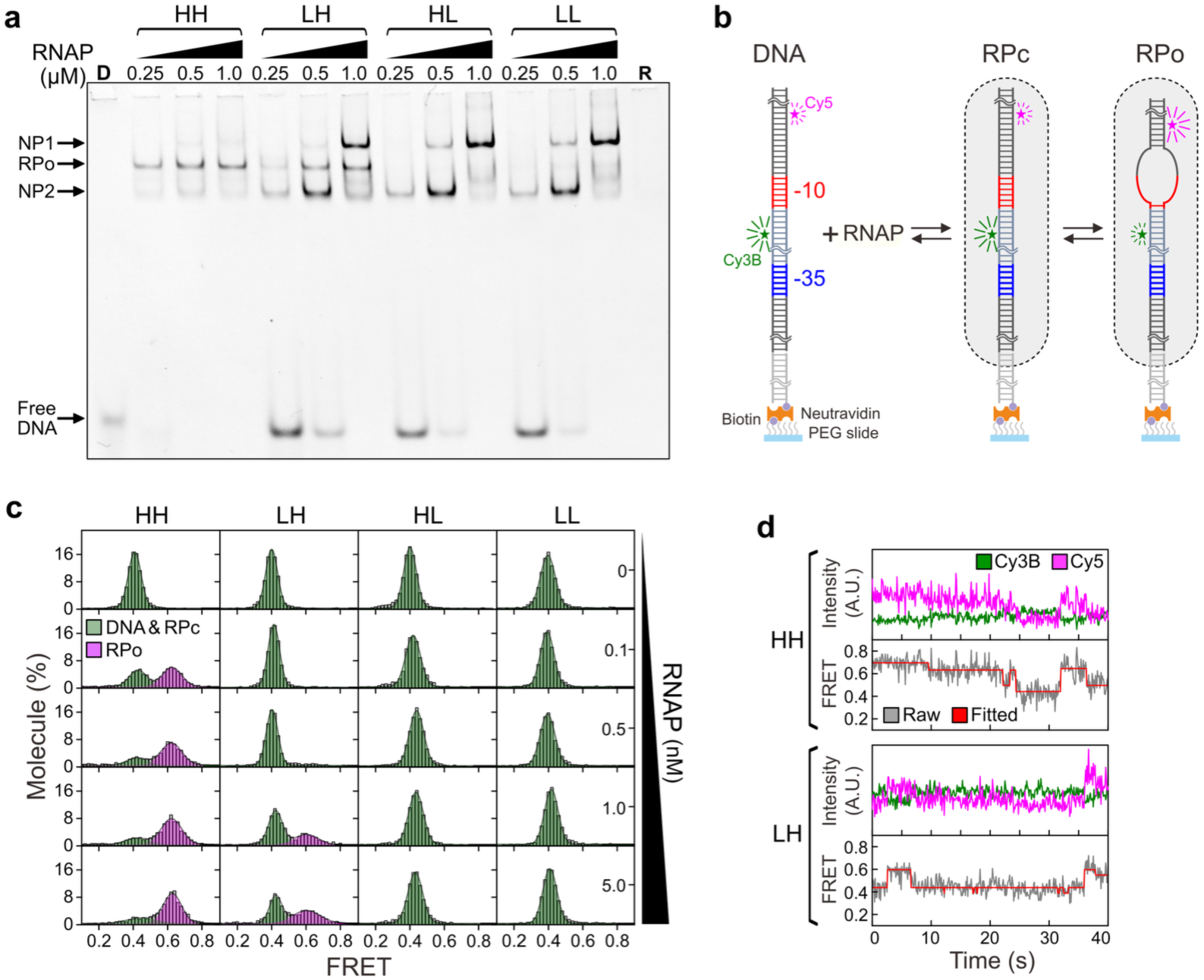
*In vitro* validation of the essentiality of −35 and −10 elements for RNAP binding. **a** Electrophoretic mobility shift assay of RNAP binding to promoters HH, LH, LL, and HL shown in Fig. 4b. Promoter DNA is supplemented at 0.25 µM throughout. D, only promoter DNA; R, only RNAP (1.0 µM); RPo, putative RNAP–promoter open complex; NP1 and NP2, non-productive binding forms 1 and 2. **b** Determining the RNAP–promoter binding kinetics by single-molecule fluorescence resonance energy transfer (FRET). Promoter DNA, attached to a polyethylene glycol (PEG)/biotin-PEG coated slide surface through biotin-Neutravidin-biotin interactions, is tagged with Cy3B and Cy5 fluorophores. RPc, RNAP–promoter close complex. **c** Equilibrium single-molecule FRET distributions across RNAP titration. Gaussian fits reveal peaks at ∼0.42 and 0.62, corresponding to the DNA-only/RPc and RPo states, respectively. Each promoter dataset includes at least 1,800 observed molecules. **d** Representative single-molecule FRET traces of the HH and LH promoters. The upper panel displays Cy3B and Cy5 fluorescence intensities, while the lower panel shows the raw FRET trajectory along with the Hidden Markov Model fit. Most traces exhibit highly dynamic transitions between the DNA-only/RPc and RPo states, frequently proceeding through intermediate states.

To clarify the EMSA observations, we performed single-molecule fluorescence resonance energy transfer (smFRET) experiments to further dissect RNAP interactions with the four promoters. LL, HL, LH, and HH promoter DNA labeled with Cy3B and Cy5 fluorophores was attached to a solid surface (Fig. 5b and Supplementary Fig. S6a). The basal FRET value (∼0.42) was comparable between free DNA and the RNAP–promoter close complex (RPc) due to the subtle change in DNA upon RNAP binding ^13^, whereas transition to the RPo state produced higher FRET signals (∼0.62) due to a conformational change that brought Cy3B and Cy5 into proximity. With increasing RNAP concentrations, smFRET experiments revealed a gradual shift of promoter populations from the DNA-only or RPc states toward the RPo state in the transcription-active HH and LH promoters but not in the transcription-silent HL and LL promoters (Fig. 5c). Notably, the RPo fraction of the HH promoter increased steeply and saturated at low RNAP concentrations (0.1–0.5 nM), whereas the LH promoter required ≥ 5 nM RNAP to saturate, with its population split evenly between the DNA-only/RPc and RPo states (Supplementary Fig. S6b), consistent with their *in vivo* expression levels and EMSA results (Fig. 4c, 5a). Even at high RNAP concentrations (5 nM), the time traces of HH and LH promoters revealed frequent interconversions between the DNA-only/RPc and RPo states (Fig. 5d), underscoring the rapid dynamics and inherent instability of the transcription bubble ^23,63,64^. Taken together, *in vivo* and *in vitro* experiments of the LL, HL, LH, and HH promoters support the “-10 essential and −35 dispensable” hypothesis of bacterial promoter function and reaffirm its audio system–like design principle, deduced from the full −35/-10 sequence−function landscapes of PL_C17_ and PL_C16_ and the promoter evolution experiment (Fig. 2g, 3a, 3b).

### Transcription activators alter the functional contribution of the −35 element

Building on findings from constitutive promoters, we next examined how the audio system–like property shapes the functional landscapes of TF-regulated promoters. We chose TetR-, LuxR-, and CueR-regulated promoters (P*_tetA_*, P*_luxI_*, and P*_copA_*, respectively) for investigation because each TF represents a distinct regulatory mode and is frequently used in synthetic biology applications (Fig. 1a and Supplementary Fig. S7a) ^65,66^. Unlike PL_C17_ that had all twelve base positions mutated, for TF-regulated promoters we selected three and four base positions in their −35 and −10 elements, respectively, for saturation mutagenesis (4^7^ = 16,384), yielding libraries PL_TetR_, PL_LuxR_, and PL_CueR_ (Fig. 6a). This mutagenesis scheme served two purposes: (1) keeping TFBSs that partially overlapped the - 35 element unaltered and (2) scaling down the library size to enhance sort-seq measurement precision. Promoter strength of PL_TetR_, PL_LuxR_, and PL_CueR_ was determined in the absence (basal) or presence (induced) of their corresponding inducers, anhydrotetracycline (aTc), acyl-homoserine lactone (AHL), and copper ions (CuSO_4_), each supplemented at a saturating concentration (Supplementary Fig. S7b). Two sort-seq replicates collectively detected 100% variants in both basal and induced conditions of the three libraries (median reads/variant = 545–1230; Supplementary Fig. S2b), with Pearson’s correlation coefficients ranged from 0.71 to 1.00 (Fig. 6b). Analysis of data passing the quality filter (≥ 20 reads and present in at least two sort-seq ranks) showed that the promoter strength distributions of the three libraries were generally lower and narrower in the basal condition while higher and wider in the induced condition (Fig. 6c; Supplementary Data S1). Promoter strength of PL_CueR_ was overall lower than that of PL_TetR_ and PL_LuxR_ in both basal and induced conditions (*P* < 10^−300^, Mann–Whitney U test), likely restrained by its overlength spacer element (19 bp) shared among MerR-type promoters ^40,41^.

**Figure 6.**
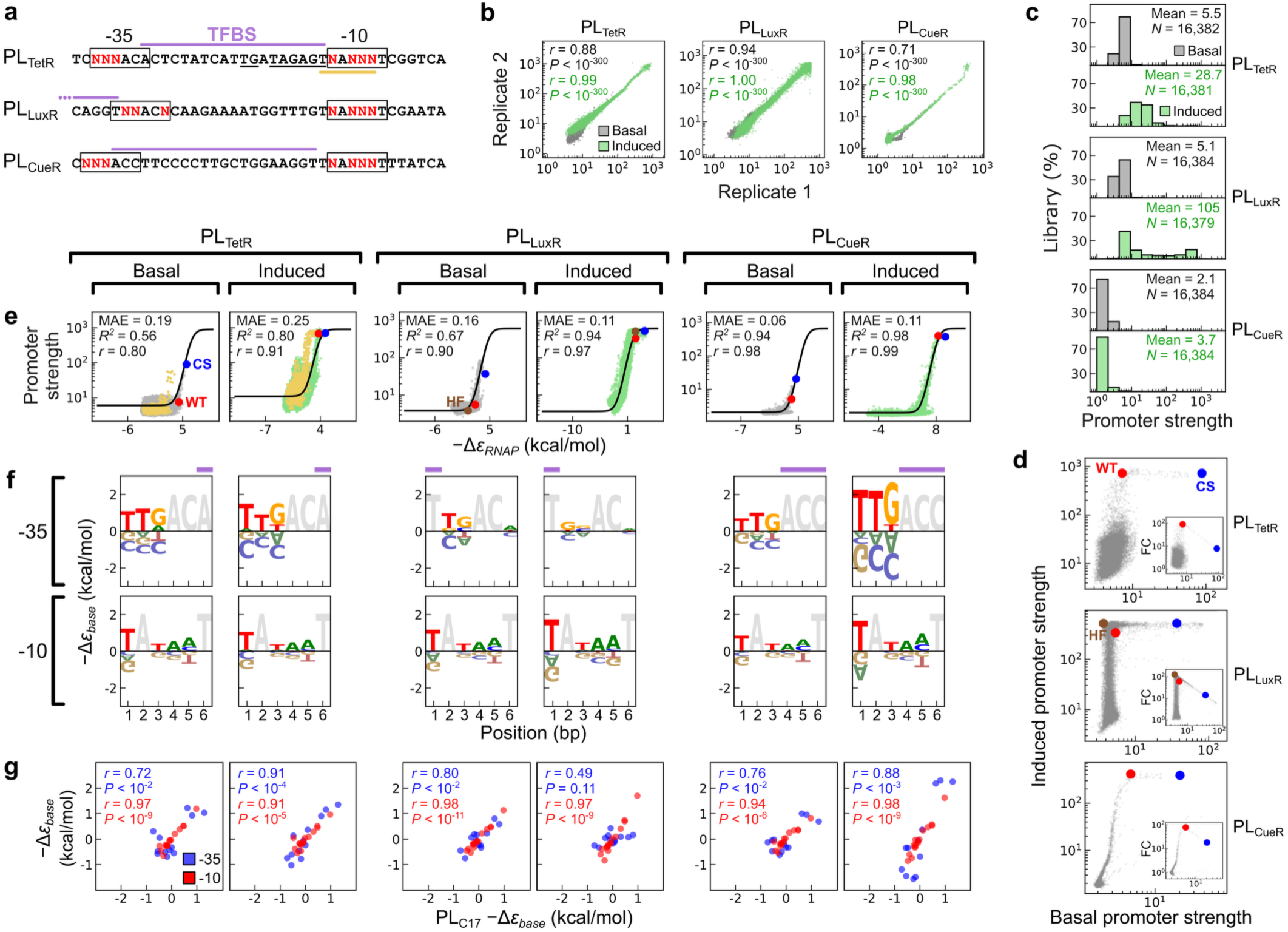
Functional contribution of the −35 and −10 elements in regulated promoters. **a** Sequence composition of the TF-regulated promoter libraries. Randomized base positions are shown in red. TFBSs are indicated by purple lines. Location of the 1-bp-shifted −10 element (TAAnnT), found in 1/16 of PL_TetR_ variants, is marked with a yellow line. The putative extended −10 and −10 motifs within the spacer element of PL_TetR_ are underlined. **b** Correlation (Pearson’s *r*, *t*-test) of promoter strength between two sort-seq replicates in the basal and induced conditions. Promoter strength is reported as the relative fluorescence unit. **c** Library distribution of promoter strength in the basal and induced conditions. **d** Correspondence between basal and induced expression levels of promoter variants. Variants carrying the consensus (CS: PL_TetR,_ TTGACA/TATAAT; PL_LuxR_, TTGACA/TATAAT; PL_CueR_, TTGACC/TATAAT) or wild-type (WT: PL_TetR_, TTGACA/TAT<u>TT</u>T; PL_LuxR_, TT<u>T</u>AC<u>G</u>/TATA<u>G</u>T; PL_CueR_, TTGACC/TA<u>ACC</u>T; underlined bases indicate differences from CS) −35/-10 sequences are highlighted. A PL_LuxR_ variant exhibiting low basal expression and a high fold change (HF, TCCACT/TATATT) is also shown. In the PL_CueR_ CS variant, the 3′ end of its −35 element is a cytosine instead of the conventional adenine to preserve its TFBS. The inset in each plot shows the correspondence between basal promoter strength and fold changes (FC) upon induction. **e** Model fitting curves of predicted binding energy (*−Δε_RNAP_*) versus promoter strength for PL_TetR_, PL_LuxR_, and PL_CueR_ variants in the basal and induced conditions. Prediction outliers in PL_TetR_, likely due to the putative extended −10 and −10 motifs shown in **a**, are marked in yellow. Variants carrying the WT, CS, or HF −35/-10 sequences are highlighted. MAE, mean absolute error. **f** Model-estimated binding energy of each nucleobase (*−Δε_base_*) within the −35 and −10 elements in the basal and induced conditions. Fixed base positions are shown in grey, with those overlapping TFBS indicated by purple lines. **g** Correlation (Pearson’s *r*, *t*-test) of *−Δε_base_* between the constitutive (PL_C17_) and TF-regulated promoters in the basal and induced conditions (*N*_-35_ = 12, *N*_-10_ = 16).

Despite differences in regulatory mechanisms, basal and induced expression of variants across the three regulated promoter libraries followed a diminishing-returns relationship (Fig. 6d): most variants had low basal expression and spanned a wide range of induced expression, whereas those with higher basal expression showed only limited expression increments upon induction. This general trend contrasts with a recent study that reported a simple inverse relationship between basal expression and the fold change (i.e., the ratio of induced to basal expression) of regulated promoters ^48^. Notably, the wild-type (WT) variants (i.e., bearing the native −35 and −10 sequences) in PL_TetR_, PL_LuxR_, and PL_CueR_ all sat at the knees of the bent distributions and exhibited high fold changes (90.9×, 59.8×, and 78.3×, respectively; Fig. 6d inset), with the library ranking as 22^nd^, 2676^th^, and 1^st^, respectively (corresponding to the top 0.13%, 16.3%, and 0%). In contrast, PL_TetR_, PL_LuxR_, and PL_CueR_ variants bearing the consensus −35 and −10 sequences (CS) all occupied the rightmost end of the plateau and exhibited lower fold changes (7.9×, 14.1×, and 18.3×, respectively), with the library ranking as 1113^th^, 4705^th^, and 103^rd^, respectively (corresponding to the top 6.8%, 28.7%, and 0.62%). The slight sequence difference (2–3 bases) between WT and CS variants suggests that WT regulated promoters are specifically selected to confer low basal expression yet high fold changes (Fig. 6d inset and legend). We then applied modeling to decipher the underlying design principles.

Models for TF-regulated promoters comprised the basal and induced parts (Supplementary Fig. S3a), fitted separately to promoter sequence–strength data from the two conditions. They differed from the PAS model (for constitutive promoters) as follows: (1) the PL_TetR_ model incorporated promoter-binding competition between the repressor TF and RNAP, reducing transcription initiation in the basal condition; (2) the PL_LuxR_ and PL_CueR_ models allowed promoter co-occupancy by activator TFs and RNAP, with activator TFs enhancing RNAP binding and transcription upon induction (represented by an energy term −*ε_act_*). Model predictions for the basal and induced expression of PL_TetR_, PL_LuxR_, and PL_CueR_ showed a sigmoidal relationship between *−Δε_RNAP_* and promoter strength that agreed well with data (*R*^2^ = 0.56*−*0.98, Pearson’s *r* = 0.80*−*0.99, MAE = 0.06−0.25; Fig. 6e). The lower prediction accuracies of the PL_TetR_-basal and PL_TetR_-induced models reflected an outlier subset (1/16 of PL_TetR_ variants, excluded from model fitting), in which the 3′ thymine of the P*_tetA_* spacer formed a 1-bp-shifted −10 element (TAAnnT; yellow in Fig. 6a, e) together with the designated −10 region. Function of this shifted −10 element was validated by mutational analysis (Supplementary Discussion and Fig. S8a). Additionally, the model-fitting curve revealed the higher background promoter activity of PL_TetR_ in the induced than in the basal condition (Fig. 6e), consistent with the presence of a shadow promoter whose extended −10 and −10 elements (TG_TAGAGT; black underline in Fig. 6a) resided in the P*_tetA_* spacer element ^67^.

Intriguingly, modeling of PL_TetR_, PL_LuxR_, and PL_CueR_ elucidated the design of TF-regulated promoters with low basal expression yet high fold changes. Moreover, it indicated that the −35 element acts the volume tuner through which activator TFs modulate the audio system−like promoter function, consistent with the conclusion from constitutive promoters. Across the three regulated promoter libraries, models showed that WT and CS variants both approached maximal induced expression (Fig. 6e), with 15.5% of PL_LuxR_ variants reaching the ceiling, yielding a bimodal promoter strength distribution (Fig. 6c). However, in the basal condition, subtle *−Δε_RNAP_* differences (1.09−3.18 kcal mol^-1^) placed WT at the foot of the sigmoidal *−Δε_RNAP_*−expression curve yet CS on its mid-slope (Fig. 6e), resulting in their marked difference in fold change (Fig. 6d inset). The results support WT promoter sequences as an evolutionary optimization: through fine-tuning *−Δε_-35_* and *−Δε_-10_*, they achieve low basal and high induced expression, thereby maximizing the dynamic range of transcriptional regulation. On the other hand, models revealed an apparent *−Δε_base_* change in the −35 element of LuxR- and CueR-regulated promoters (Fig. 6f). Relative to constitutive promoters, the magnitude of *−Δε_base_* in the −35 element decreased in PL_LuxR_ but increased in PL_CueR_ upon induction, whereas *−Δε_base_* in the −10 element remained comparable across constitutive promoters and both basal and induced conditions of the three regulated libraries (Fig. 6g). These findings provide the first quantitative evidence, showing that activator TFs regulate transcription by specifically modulating the −35 element’s binding energy contribution, thereby extending previous biochemical characterizations ^38,45,49^. This conclusion was further supported when each regulated promoter model was refitted under the stricter constraint of using a single −Δ*ε_base_* set for both basal and induced conditions (Supplementary Fig. S8b−d; see Methods).

To examine the generality of our conclusion from PL_TetR_, PL_LuxR_, and PL_CueR_, we divided *E. coli* σ^70^ promoters into constitutive (i.e., lacking RegulonDB-annotated TFBS) and TF-regulated classes, the latter further subclassified by regulatory mode ^68^. We then applied the PAS model to identify the −35 and −10 elements in each promoter and compute *−Δε_-35_* and *−Δε_-10_*. Leveraging availability of the dataset ^28^, this analysis also included promoters regulated by MerR-type TFs from diverse bacteria. Consistent with prior knowledge and findings in PL_TetR_, PL_LuxR_, and PL_CueR_ ^2,26^, only Class II and MerR-type promoters showed significantly different *−Δε_-35_* relative to constitutive promoters (*P* < 0.001, Mann–Whitney U test), with *−Δε_-10_* of MerR-type promoters significantly lower than that of constitutive promoters (*P* < 0.001; Fig. 3c). In sum, the results underscore the significance of core promoter elements in transcriptional regulation, elucidate the design principles underlying their evolutionary optimization to attain high fold changes, and quantitatively demonstrate that activator TFs target the −35 element to tune promoter strength.

### Leaky expression of P*_luxI_* is critical for quorum sensing

Although WT variants in PL_TetR_, PL_LuxR_, and PL_CueR_ all exhibited high fold changes, PL_LuxR_ WT had a poorer library ranking (2676th vs. 22nd and 1st), reflected by its position at the inner rather than outer corner of the basal expression–fold change distribution (Fig. 6d inset). Why would evolution favor the P*_luxI_* sequence (i.e., PL_LuxR_ WT) that gives slight basal expression at the expense of fold change? In its native context, P*_luxI_* controls expression of LuxI (AHL synthase), which produces the inducer for LuxR, forming a positive feedback loop to activate P*_luxI_* expression ^69^. We hypothesized that this leaky expression may establish a critical baseline for initiating quorum sensing—a mechanism for receiving and responding to AHL produced by neighboring cells ^69^. To test the hypothesis, we generated a P*_luxI_*-*gfp*-*luxI* cassette (*luxI*(*+*)) by fusing the *luxI* gene, including its native ribosome binding site, downstream of the P*_luxI_*-*gfp* cassette (*luxI*(*−*); Fig. 7a). From each cassette, we created three variants whose P*_luxI_* −35 and −10 sequences matched WT, CS, and a high-fold-change (HF) variant that exhibited minimal basal expression and a 126.4-fold increase upon induction (ranked 15^th^, top 0.09% in PL_LuxR_; Fig. 6d). Characterization of the three *luxI*(*−*) variants showed that their promoter strength increased with exogenously supplied AHL and attained similar maximal expression levels in liquid culture (Fig. 7b). HF.*luxI*(*−*), WT.*luxI*(*−*), and CS.*luxI*(*−*) exhibited minimal, slight, and significant basal expression (*i.e.*, AHL = 0 μM) during the exponential phase, respectively, consistent with the sort-seq result and model-inferred *−Δε_RNAP_* (Fig. 6d, e).

**Figure 7.**
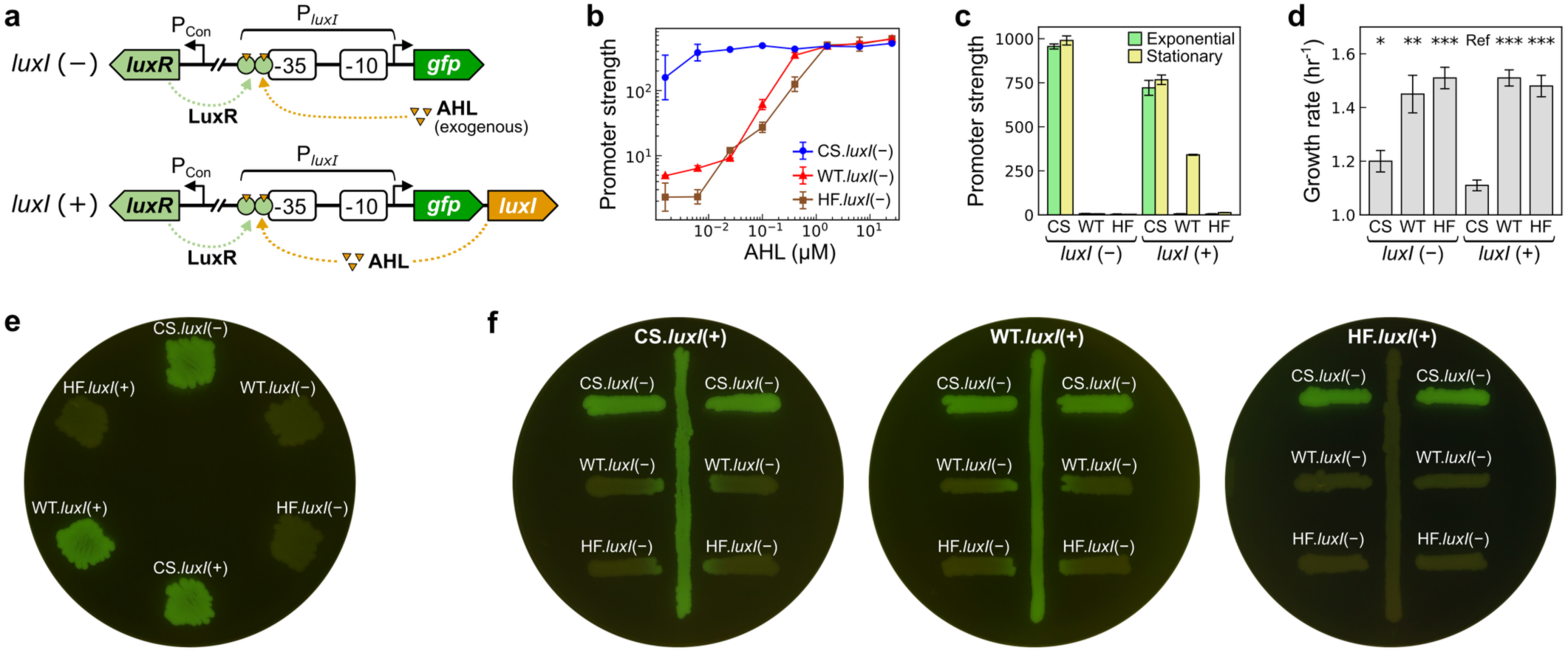
Influence of the −35 and −10 sequences on P*_luxI_* expression and quorum sensing in *E. coli*. **a** P*_luxI_* expression cassettes lacking (*−*) or containing (+) the acyl-homoserine lactone (AHL)-producing gene *luxI*. The P*_luxI_* −35/-10 sequences in the *luxI*(*−*) and *luxI*(*+*) cassettes are replaced by those of three PL_LuxR_ variants—consensus (CS, TTGACA/TATAAT), wild-type (WT, TTTACG/TATAGT), and high-fold-change (HF, TCCACT/TATATT; Fig. 6d)—to generate six cassette variants. **b** Promoter strength of the *luxI*(*−*) variants in response to exogenous AHL supplementation in the exponential-phase liquid culture. **c** Promoter strength of the *luxI*(*−*) and *luxI*(*+*) variants in liquid culture without AHL supplementation at the exponential and stationary phases. **d** Growth rate of the *luxI*(*−*) and *luxI*(*+*) variants in the exponential-phase liquid culture without AHL supplementation. CS.*luxI*(*+*) serves as the reference (Ref) for statistical comparisons. Statistical significance from one-way ANOVA with Dunnett’s post-hoc test is indicated as follows: \*\*\**P* < 0.001; \*\**P* < 0.01; \**P* < 0.05. **e** Promoter strength of the *luxI*(*−*) and *luxI*(*+*) variants on an agar plate without AHL supplementation. **f** Quorum sensing assay for AHL production and perception of *luxI*(*+*) and *luxI*(*−*) variants, respectively, on agar plates without AHL supplementation. In **b**, **c**, **e**, **f**, P*_luxI_* expression is reported by cellular GFP expression. In **b**–**d**, data are presented as means and standard deviations from three independent measurements.

Upon quantifying promoter strength of the *luxI*(*−*) and *luxI*(*+*) variants in liquid culture without exogenously supplemented AHL, we found that CS.*luxI*(*−*) and CS.*luxI*(*+*) exhibited strong expression across growth phases, HF.*luxI*(*−*), HF.*luxI*(*+*) and WT.*luxI*(*−*) showed minimal expression throughout, and WT.*luxI*(*+*) yielded significant expression only after entering the stationary phase (Fig. 7c). The results highlight the importance of leaky WT P*_luxI_* expression in cell density-dependent quorum sensing. Notably, promoter strength of CS.*luxI*(*+*) was significantly lower than that of CS.*luxI*(*−*) in the absence of exogenous AHL (*P* < 0.01, *t*-test). This phenomenon may result from the growth burden imposed by high expression of LuxI in addition to GFP in CS.*luxI*(*+*), as reflected by its significantly lower growth rate compared with other variants (Fig. 7d; Dunnett’s test, *P* < 0.05).

To assess whether the three *luxI*(*+*) variants produce sufficient AHL to trigger quorum sensing at a distance, we cultured them and the three *luxI*(*−*) variants on agar plates without exogenous AHL. Similar to the stationary-phase liquid culture (Fig. 7c), the HF.*luxI*(*+*) lawn was dim while WT.*luxI*(*+*) and CS.*luxI*(*+*) emitted significant GFP fluorescence (Fig. 7e). When growing *luxI*(*+*) and *luxI*(*−*) variants on the same plate, we found that both WT.*luxI*(*+*) and CS.*luxI*(*+*) induced GFP expression of HF.*luxI*(*−*) and WT.*luxI*(*−*) in proximity, whereas HF.*luxI*(*+*) could not (Fig. 7f). The CS.*luxI*(*−*) lawn emitted significant GFP fluorescence even when co-culturing with the poor AHL-producing HF.*luxI*(*+*) variant, indicating that the strong basal expression of CS.*luxI*(*−*) rendered it insensitive to AHL sent from neighboring cells. With HF.*luxI*(*+*) producing insufficient AHL and CS.*luxI*(*−*) unresponsive to surrounding signals, results support the leaky expression of WT P*_luxI_* as an evolutionary optimization for quorum sensing.

## Discussion

Previously, we applied the PAS model to map the promoter architecture upstream of genome-wide transcription start sites across 49 diverse bacteria ^11^. Consistent with findings in model species ^14,70–72^, the −10 and −35 elements exhibited the highest and the second-highest sequence conservation, respectively, among core promoter elements. To assess their roles in promoter birth and transcriptional regulation, here we comprehensively determined the −35 and −10 sequence–function landscapes in constitutive and TF-regulated promoters by high-throughput experiments. Through quantitatively dissecting their contributions to constitutive promoter strength and promoter birth in laboratory evolution (Fig. 2, 3), followed by *in vivo* and *in vitro* experimental validations (Fig. 4, 5), we showed the −10 element to be essential whereas the −35 element is dispensable for promoter identity from a genomic and evolutionary perspective. The results corroborate prior biochemical characterizations of the RPo complex, which emphasize the essentiality of the −10 element for promoter melting and maintaining structural stability ^73–77^. The central role of the −10 element suggests that bacterial promoter prediction should begin by identifying the −10-like motif within DNA fragments ^72^, whereas undesired transcriptional activity within coding sequences can be effectively reduced by excluding −10-like motifs during gene synthesis ^78^.

Contrary to the ON/OFF switch role of the −10 element in promoter identity, the greater −35 sequence variation across the domain Bacteria could be attributed to its minor importance in this aspect (Fig. 2–5) ^11^, and most interestingly, to its diverse interactions with RNAP and TFs (Fig. 6). As reviewed previously ^2,27,66^, the −35 element often overlaps with the binding sites of activator TFs that function via DNA bending (MerR family) or allosteric interaction with σ^70^_4_ (Class II; Fig. 1a). By systematically characterizing the −35 and −10 sequences in promoters regulated by representative TFs, CueR (MerR family) and LuxR (Class II), our results provide genetic and mechanistic evidence that illustrates the coevolution of core promoters, RNAP, and TFs. In both cases, the WT variants exhibited low basal and high induced expression, placing their fold changes among the highest in their respective libraries (Fig. 6d inset), indicative of adaptive evolution. The peaked tradeoff relationship between basal expression and fold change in PL_CueR_, PL_LuxR_, and PL_TetR_ (16,384 variants each; Fig. 6d inset) contrast with a simple inverse relationship reported in a recent study that examined only 153 −35 sequence variants per TF ^48^. The distinct data distributions likely reflect a substantial difference in sampling coverage, suggesting that the provocative idea proposed therein—repressor TFs inhibit transcription by over-stabilizing rather than blocking RNAP–promoter binding—deserves careful reconsideration. Beyond this, the bent basal–induced expression distributions of our TF-regulated promoter libraries revealed numerous low-basal-expression variants with a wide range of induced expression (Fig. 6d), underscoring the core promoter as a key dimension for engineering transcriptional regulation and highlighting the utility of our datasets for illuminating the design principles for synthetic biology applications (Supplementary Data S1). The necessity of leaky P*_luxI_* expression for LuxR-regulated quorum sensing provides a compelling example (Fig. 7).

Notably, model analyses of the PL_CueR_ and PL_LuxR_ libraries revealed significant increases and decreases in *−Δε_-35_*, respectively, but negligible changes in *−Δε_-10_* under the induced condition (Fig. 6f, g). Rather than modeling artifacts, a recent cryo-EM study shows that CueR and σ^70^_4_ form a sandwich clamp that tightly grasps the −35 element from the opposite side of the DNA helix ^45^, indicating that the elevated *−Δε_-35_* in PL_CueR_ represents a reliable quantitative measure complementary to the biochemical evidence. On the other hand, the LuxR–promoter–RNAP complex structure is unavailable for comparison owing to inherent difficulties in purifying the LuxR protein ^79^. The reduced *−Δε_-35_* of PL_LuxR_ upon LuxR binding suggests a more extensive overlap between its TFBS and the - 35 element than is currently recognized (Fig. 6a) ^49^. If true, this would identify LuxR as an ambidextrous TF—similar to SoxS, Rob, and MarA ^80^—that functions by occluding σ^70^_4_ from the −35 element in addition to its previously known physical contacts with σ^70^_4_ and αCTD ^38,81^. Together, these findings reaffirm the tuner role of the −35 element in bacterial promoter function and demonstrate the power of our high-throughput genetics/modeling framework to resolve transcriptional mechanisms and protein–DNA interactions otherwise inaccessible to conventional biochemical approaches.

## Methods

### Strains and growth media

*E. coli* MG1655 (wild type) served as the host strain for most investigations, while its metal homeostasis-compromised derivative, EK317 (Δ*cueR*, Δ*copA*, Δ*cusCFBA*, Δ*zntR*, Δ*zntA*), was used to assay PL_CueR_ ^84^. LB medium consisted of 10 g NaCl (Fisher Scientific), 10 g bacterial tryptone (BD Bacto), and 5 g yeast extract (BD Bacto) in one liter of deionized water. LBK medium, LB medium supplemented with filter-sterilized kanamycin sulfate (50 mg/L, Thermo Fisher Scientific), was used for growing cells for plasmid extraction. LBKG medium, LBK medium supplemented with filter-sterilized glucose (10 g/L, Sigma-Aldrich) to enhance GFP expression, was used for growth assays or fluorescence-activated cell sorting (FACS) experiments. LBKGE medium, LBKG medium supplemented with filter-sterilized ethylenediaminetetraacetic acid (EDTA; 50 μM, Riedel-de Haën) to remove residual metal ions, was used for culturing *E. coli* EK317 bearing promoter P*_copA_* and its variants ^85^.

### Plasmid construction

All primers and oligonucleotides were synthesized by Integrated DNA Technologies and described in Supplementary Table S1. All plasmids were listed in Supplementary Table S2. Most plasmids were constructed by inverse PCR, while some were made by restriction digestion and ligation. Primers used in inverse PCR were 5’ phosphorylated using T4 polynucleotide kinase (NEB). Inverse PCR was conducted using Phusion DNA polymerase (Thermo Fisher Scientific). DpnI restriction enzyme (NEB) was employed to remove template plasmids. T4 DNA ligase (Thermo Fisher Scientific) was utilized to circularize plasmids. After purification by the Zymo DNA Clean & Concentrator kit, the ligation product was transformed into *E. coli* using the BIO-RAD MicroPulser electroporator. Electrocompetent cells were prepared following an established protocol ^86^.

Plasmids pJU84 and pWS24, bearing P*_purR_* and a synthetic promoter P*_arti_*, respectively, were constructed previously ^11^ and served as templates for synthesizing constitutive promoter libraries PL_C17_ and PL_C16_. Template plasmids pTK77, pTK82, and pTK76 for regulated promoter libraries PL_TetR_, PL_LuxR_, and PL_CueR_, respectively, were constructed through the following steps. First, *tetR*, *luxR*, and *cueR* genes from pRF-TetR, pLuxR-RP, and pYK01, respectively ^65,87,88^, were inserted into pWS24 by restriction and ligation, leading to three intermediate plasmids, pTK59, pTK63, and pTK47. They further went through tree inverse PCR modifications to generate pTK77, pTK82, and pTK76: (1) introducing a constitutive promoter (P*_cueR_*) to drive TF gene expression ^88^; (2) introducing TF-regulated promoters P*_tetA_*, P*_luxI_*, and P*_copA_*, respectively, to control *gfp* gene expression; (3) tuning the ribosome-binding site (RBS) for a suitable dynamic range of GFP expression while minimizing growth inhibition from expression burden (Supplementary Fig. S7a).

We performed the Golden Gate assembly to introduce the *luxI* gene from *Vibrio fischeri* ATCC7744 downstream of the P*_luxI_*-controlled *gfp* gene in plasmids pTK82, pWS51, pML14, resulting in pML85, pML86, pML87, respectively. The *luxI* gene fragment was amplified by PCR using the primer pair GIp1/GIp2, and the vector backbones of pML85, pML86, pML87 were amplified using GVp1/GVp2. The *luxI* and vector fragments were pooled together in a 3:1 molar ratio, digested with Eco31I (Thermo Fisher Scientific), and ligated with T4 DNA ligase following the standard incubation conditions ^89^.

### Construction of promoter libraries

Promoter libraries PL_C17_, PL_TetR_, PL_LuxR_, and PL_CueR_ were constructed by inverse PCR using pJU84, pTK77, pTK82, and pTK76 as template plasmids, respectively, Q5 DNA polymerase (NEB), and primers pre-phosphorylated and HPLC-purified by Integrated DNA Technologies. The PL_C16_ library was generated via the same procedure, except that it was constructed and assayed as four complementary subsets (PL_C16-1_ to PL_C16-4_). Following DpnI treatment, T4 DNA Ligase was utilized to circularize plasmids. After DNA purification, ligation products were mixed with 180 µL of freshly made electrocompetent cells and divided into six equal aliquots for electroporation. All transformed cells were pooled and revived in 6 mL of Recovery Medium at 37°C and 225 rpm for 1 hour, transferred to 30 mL of LBK medium, and incubated at 37°C and 225 rpm overnight. The transformation efficiency for library construction was ≥ 5×10^7^ colony forming units. Cells in the overnight culture were collected by centrifugation, resuspended in 1 mL of PBS containing 25% glycerol (v/v), aliquoted into 20 portions, and stored at *−*80°C as the inoculum for sort-seq experiments.

### Characterization of cell growth, GFP expression, and quorum sensing

Single-cell fluorescence was quantified by a BD FACSJazz cell sorter. Each experiment began with inoculation of 1 μL of frozen stocks into 2 ml of LBKG (most promoters) or LBKGE (P*_copA_* and variants) media incubated at 37°C and 225 rpm overnight. Next day, 5 μL of the pre-culture was transferred into 2 mL of LBKG medium, with or without inducers. Three inducers were supplied at saturating concentrations (Supplementary Fig. S7b): anhydrotetracycline hydrochloride (aTc, 100 μg/L, Alfa Aesar) for TetR, N-(β-ketocaproyl)-L-homoserine lactone (AHL, 10 μM, Sigma-Aldrich) for LuxR, and CuSO_4_·5H_2_O (10 μM, J. T. Baker) for CueR. When OD_600_ reached 0.55–0.65 at 1 cm path length, 100 μl of this culture was mixed thoroughly with 900 μl of pre-chilled phosphate-buffered saline (PBS) and was stored on ice. For each strain, the cell sorter quantified the GFP fluorescence (excitation / emission: 488 nm / 513 ± 8.5 nm) of 50,000 to 200,000 cells in three replicate cultures. For the quorum-sensing assay, *E. coli* bearing the *luxI*(−) and *luxI*(+) cassette variants were grown in LBK medium at 37 °C and 225 rpm overnight and then transferred onto LBK agar plates using an inoculating loop. After incubation at 37 °C overnight, GFP fluorescence from bacterial lawns was visualized using a BluView Transilluminator (MBE-200A). Images were captured with a digital camera (ISO 125, shutter speed 1/20 s, color temperature 3300 K).

### RNA extraction and quantitative PCR

*E. coli* was grown in 10 ml of LBKG medium at 37°C and 225 rpm overnight. When OD reached 0.55–0.65, cells were harvested by adding 1/10 the volume of a growth-stopping solution (5% Tris-EDTA saturated phenol and 95% ethanol), followed by centrifugation at 9000 g for 10 minutes at 4°C. Total RNA was extracted using the RNeasy Mini Kit (QIAGEN), followed by the removal of residual genomic DNA with the Turbo DNA-free Kit (Thermo Fisher Scientific). cDNA was synthesized by the High-Capacity cDNA Reverse Transcription Kit (Thermo Fisher Scientific). For each variant, mRNA extraction and cDNA synthesis were performed twice independently. The primer pairs for detecting the *gfp* and *gapA* transcripts were YLEp5/YLEp6 and YLEp7/YLEp8, respectively. Real-time PCR of each cDNA sample was performed in two replicates with the iQ SYBR Green Supermix (Bio-Rad) on a CFX Connect Real-Time PCR System (Bio-Rad). Only measurements with a standard deviation of cycle threshold values ≤ 0.2 are retained for subsequent analysis. The *gapA* housekeeping gene was chosen as the reference for data normalization. mRNA levels were calculated based on an established method ^90,91^.

### Sort-seq experiment

All promoter libraries, except PL_CueR_, were revived by inoculation of one aliquot of frozen stocks into 20 mL of LBKG medium. PL_CueR_ was instead revived in LBKGE medium to remove residual metal ions ^84^. Meanwhile, spike-in variants, promoters with known sequences and expression levels (Supplementary Table S3), were revived by inoculation of 1 μL of frozen stocks into 1 mL LBKG medium. Promoter libraries and spike-in variants were grown overnight at 37°C and 225 rpm. Subsequently, 50 μL and 2.5 μL of the library and spike-in precultures were transferred to 20 mL and 1 mL LBKG medium, respectively. Both cultures were grown at 37°C and 225 rpm until their OD reached 0.55–0.65. All spike-in variants were evenly pooled together and added into the library culture in a 1:2500 ratio (v/v). This mixture was diluted 10-fold with PBS and stored on ice. FACS was performed by a BD FACSJazz cell sorter, and cells were maintained at 4°C throughout. The sensitivity of the photomultiplier tube was calibrated based on pWS19 and pYC08 ^86^. The fluorescence distribution of each library was divided into 6–8 ranks, numbered in the ascending order according to the fluorescence intensity. A total of 1.5×10^7^ and 5×10^6^ cells were collected for constitutive and regulated promoter libraries, respectively, with the number of cells collected for each rank proportional to its relative abundance in the library (Supplementary Table S4). After FACS, cells in each rank were immediately grown in 10 ml LBK medium at 37°C and 225 rpm overnight. Subsequently, the plasmids of each rank were extracted by the Qiagen Plasmid Miniprep Kit. To inspect the FACS purity of each rank, 5 μL of overnight cultures were inoculated into 2 mL of LBKG medium with or without inducers incubated at 37°C and 225 rpm. The fluorescence distribution was examined by the cell sorter when OD_600_ of the culture reached 0.55–0.65 (Supplementary Fig. S1). Promoter sequences in each sorting rank were determined by the Illumina NovaSeq 6000 system. Sequencing library preparation followed the Illumina two-step PCR procedure. A 0.3-kb fragment, including the promoter, 5′ untranslated region, and partial *gfp* gene, was amplified by amplicon PCR using KAPA HiFi DNA polymerase (Roche), along with equal-volume mixed forward primers ILp11, ILp11a, ILp11b, and ILp11c and the reverse primer ILp12. Amplicon PCR products were purified by AMPure XP Beads (Beckman) and then subjected to index PCR using KAPA HiFi DNA polymerase and a unique pair of index primers from the Nextera XT Index Kit (Illumina). Index PCR products were purified by AMPure XP Beads, whose DNA concentrations were measured by a Qubit fluorometer (Thermo Fisher Scientific). Index PCR products were pooled based on the proportion of each rank in the library. The multiplexed sample was subjected to 150-bp paired-end sequencing. Sequencing of the constitutive and regulated libraries generated approximately 75 million and 25 million reads per experiment, respectively. We set FastQC quality score Q ≥ 25 to filter out poor sequencing results.

### Estimating promoter strength from sort-seq data

Sequence hard-matching removed irrelevant or indel-containing reads from the raw sequencing data. The sequence composition of the −35 and −10 elements was extracted from each read, and the occurrence of each promoter variant was counted in each sort-seq rank. Let *F_v_*(*n*) denote the frequency of promoter variant *v* in rank *n*, where ∑_*n*_ *F_v_*(*n*) = 1. The rank mean *R_v_* of variant *v* was calculated as *R_v_* = ∑_*n*_*n* · *F_v_*(*n*). Following identical procedures, the rank means of spike-in variants were computed. The relationship between *R_v_* and *E_v_* (i.e., GFP fluorescence on a log_10_ scale) was described as *E_v_* = *a* · *R_v_* + *b*. Linear regression of the rank means of spike-in variants and their GFP expression measured by flow cytometry was performed (Supplementary Table S5). The resulting linear equation was applied to converting the rank mean of a promoter variant into its promoter strength.

### Model construction

Models are built upon prior theoretical frameworks ^15,35,61^. They assume that promoters exist in multiple states, in terms of their interactions with RNAP and TFs. Each state of promoter variants in a library was assigned a relative probability based on its Boltzmann weight *e^-βΔε^*, where −Δ*ε* denotes the free energy, *β =* 1/*RT*, gas constant *R* = 1.98e-3 kcal mol^-1^ K^-1^, and *T* = 310 K (Supplementary Fig. S3). The models define the transcription rate as *r_max_* when RNAP is bound to the promoter, whereas a basal measurement noise *r_min_* exists in the remaining states, with *r_min_* ≪ *r_max_*. Promoter strength is calculated as the probability-weighted sum of transcription across states, where sets *B* and *U* refer to promoter states bound or unbound to RNAP, respectively:

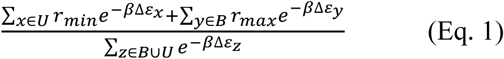

We adopted the Promoter Architecture Scanning (PAS) model from our previous work to analyze the constitutive promoter libraries, PL_C17_ and PL_C16_ ^11^. It assumed that promoters exist in unbound or RNAP-bound states (Supplementary Fig. S3), with the free energies of the two states defined as 0 and −Δ*ε_RNAP_*, respectively. −Δ*ε_RNAP_* comprises the free energies of the −35 element (−Δ*ε_-35_*), −10 element (−Δ*ε_-10_*), spacer element (*−Δε_Spacer_*), and promoter context (−Δ*ε_0_*). Each base in the −35 and −10 elements (−Δ*ε_base_*) contributes additively to −Δ*ε_RNAP_*, and −Δ*ε_Spacer_*(*l*) varies with the spacer length *l* regardless of sequence ^8^. During model fitting, each promoter variant was scanned to identify the −35 element, −10 element, and spacer length (16–18 bp) that maximize −Δ*ε_RNAP_*. Constitutive promoter strength is computed as:

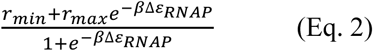

The models for TF-regulated promoter libraries comprise 3–4 molecular states differing in RNAP-promoter-TF interactions (Supplementary Fig. S3). PL_TetR_ exists in three states (free energy terms defined in parenthesis): unbound (0), RNAP-bound (−Δ*ε_RNAP_*), or TetR-bound (−Δ*ε_TetR_*). Among these, the TetR-bound and RNAP-bound states dominate under the basal condition, whereas the unbound and RNAP-bound states dominate under the induced condition. PL_LuxR_ exist in four states: unbound (0), RNAP-bound (−Δ*ε_RNAP_*), LuxR-bound (−Δ*ε_LuxR_*), or RNAP–LuxR-bound (–(Δ*ε_RNAP_* + Δ*ε_LuxR_* + *ε_act_*)), where *−ε_act_* denotes the free energy contributed by the LuxR–RNAP interaction ^81,38^. Among these, the unbound and RNAP-bound states dominate under the basal condition, whereas the LuxR-bound and RNAP–LuxR-bound states dominate under the induced condition. PL_CueR_ exists in four states: CueR^apo^-bound (−Δ*ε_CueR_apo_*), RNAP–CueR^apo^-bound (–(Δ*ε_RNAP_* + Δ*ε_CueR_apo_*)), CueR^Cu^-bound (−Δ*ε_CueR_Cu_*), or RNAP–CueR^Cu^-bound (–(Δ*ε_RNAP_* + Δ*ε_CueR_Cu_* + *ε_act_*)). CueR^apo^ and CueR^Cu^ denote CueR lacking or containing the copper ion, respectively, and −*ε_act_* denotes the free energy contributed by the CueR^Cu^-mediated DNA conformational change ^45^. Among these, the CueR^apo^-bound and RNAP–CueR^apo^-bound states dominate under the basal condition, whereas the CueR^Cu^-bound and RNAP–CueR^Cu^-bound states dominate under the induced condition. Promoter strength of PL_TetR_, PL_LuxR_, and PL_CueR_ under the basal and induced conditions is given by Eq. 3 and Eq. 4, respectively, where −Δ*ε_Spacer_* is omitted because it remains constant due to the fixed positions of the −35 and −10 elements in these libraries:

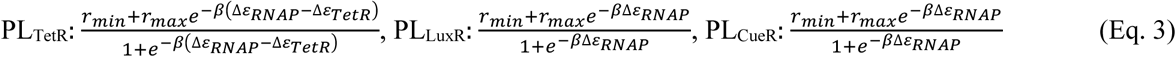

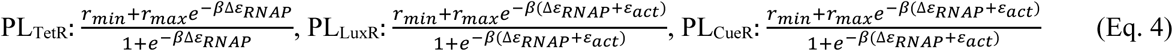

### Model fitting

We divided the promoter strength distributions of constitutive and regulated promoter libraries into 10 and 8 equal-sized bins, respectively, and randomly sampled 2,000 and 1,000 variants per bin (20,000 and 8,000 variants in total) for model fitting ^60^. We then fitted each model to its corresponding library dataset using PyTorch’s Adam optimizer to minimize the mean squared error ^92^. Model fitting followed the framework of our previous study ^11^, with the fitting algorithm replaced by Adam-based optimization to improve model performance. The performance was evaluated using the coefficient of determination (*R*^2^), the Pearson’s correlation coefficient (*r*), and the mean absolute error (MAE), computed from evenly sampled variants across each library’s promoter-strength distribution (Fig. 2d, 6e and Supplementary Fig. S8b).

Model fitting estimated the energy contribution of each base (*−Δε_base_*) in the −35 and −10 elements (Fig. 2e, 6f and Supplementary Fig. S8c), as well as other parameters (Supplementary Table S6). When training the PAS model on the datasets from two constitutive promoter libraries, we set *−Δε_Spacer_*(17) = 0 to avoid parameter degeneracy ^11^. We employed two fitting approaches for the models of TF-regulated promoter libraries, each comprising two parts describing the basal and induced conditions (Supplementary Fig. S3a). In the first approach, the basal and induced parts were fitted independently to their respective datasets, resulting in two distinct parameter sets. Because the scarcity of high-expression variants in the basal condition rendered estimation of its *r_max_* infeasible, the basal *r_max_* was set equal to *r_max_* in the induced condition (Fig. 6e). We tested parameter sensitivity by performing model fitting 30 times with *r_max_* treated as a free parameter in the basal condition and found that model-inferred *−Δε_base_* values were similar to those obtained from *r_max_*-fixed models (Pearson’s *r* = 0.98–0.99 across three libraries). Similarly, because independent estimation of *−Δε_0_* with −Δ*ε_TetR_* in PL_TetR_ (basal) and *−Δε_0_* with *−ε_act_* in PL_LuxR_ and PL_CueR_ (induced) was infeasible, we set −Δ*ε_TetR_* and *−ε_act_* to zero to avoid parameter degeneracy. We tested parameter sensitivity by performing model fitting 30 times with −Δ*ε_TetR_* and *−ε_act_* treated as free parameters and found that model-inferred *−Δε_base_* values were similar to those obtained from models with −Δ*ε_TetR_* and *−ε_act_* fixed to zero (Pearson’s *r* = 0.98–1.00). In the second approach, the basal and induced parts were fitted to their respective datasets but constrained to share a single common parameter set. An exception was made for PL_TetR_, where *r_min_* was separately estimated for its basal and induced conditions. The higher *r_min_* upon induction likely reflected additional expression from a shadow promoter (black underline in Fig. 6a), as evidenced by the elevated lower bound of its model-fitting curve relative to the basal condition (Fig. 6e and Supplementary Fig. S8b).

### Model prediction of the −35 and −10 elements in native and experimentally evolved promoters

We applied the PAS model, trained on the PL_C17_ dataset, to predict the −35 and −10 elements in *E. coli* σ^70^ promoters ^68^, in promoters regulated by the MerR family from 38 species ^28,83^, and in promoters from an experimental evolution study ^7^. For *E. coli* σ^70^ promoters, the model scanned positions −60 to +21 relative to the transcription start site (+1) to identify the −35 and −10 elements. Further, we classified them into constitutive (defined by the absence of annotated TFBSs in RegulonDB ^68^) or five TF-regulated types: (1) steric hindrance—repressor binding sites located from positions −60 to +10; (2) roadblock—repressor binding sites located from +11 to +60 ^75^; (3) deformation, including one LacI- and six GalR-regulated promoters; (4) Class I activation—activator binding sites located from −100 to −50; (5) Class II activation—activator binding sites located from −49 to −35. During classification, a promoter with more than one TFBS was assigned to multiple TF-regulated types. For promoters from the evolution study, we excluded cases where adaptive mutations occurred outside the 103-bp designated promoter region, ultimately retaining 17 pairs before and after evolution for analysis.

### RNAP purification and electrophoretic mobility shift assay

The *E. coli* RNAP holoenzyme was prepared by expressing the core enzyme and σ^70^ using plasmids pEcrpoABC(-XH)Z, pACYCDuet-LEc-rpoZ (both from the Darst lab, Rockefeller University), and pET21b-RpoD, respectively, in *E. coli* BL21, followed by purification and reconstitution using established protocols (Supplementary Table S2)^93^. DNA duplexes (59 bp) corresponding to promoter variants HH, LH, HL, and LL were prepared by annealing oligonucleotide pairs HH_F/HH_R, LH_F/LH_R, HL_F/HL_R, and LL_F/LL_R (Supplementary Table S1), respectively, at 200 μM in 10 mM HEPES (pH 8.0), 100 mM NaCl, and 1 mM MgCl_2_, using a standard melting and gradual cooling protocol (heating to 95°C, followed by cooling at 1°C per minute to 25°C). Each binding reaction was assembled in a total volume of 10 μl containing 250 nM promoter DNA and 0.01–2 μM RNAP in binding buffer (10 mM Tris-HCl, pH 8.0, 5% [v/v] glycerol, 0.1 mM EDTA, 200 mM NaCl, and 1 mM DTT). Following incubation at room temperature for 1 h, samples were resolved on 5% native polyacrylamide gels in 0.5× TBE at 100 V, and DNA-protein complexes were visualized under UV illumination after ethidium bromide staining.

### Single-molecule fluorescence resonance energy transfer assay

The non-template strand containing a 28-nt handle DNA and the template strand, each carrying an NH₂–C₆ linker at designated labeling sites (Supplementary Fig. 6a), were synthesized and HPLC-purified by Genomics. Strands were diluted to 35 µL ∼50 µM in 100 mM sodium tetraborate buffer (pH 8.4) and reacted with 35 µL 8.7 mM (∼180 equivalent) Cy3B or sulfo-Cy5 NHS ester (Lumiprobe, USA) by gentle rotation at room temperature for 24 h in an Eppendorf tube. Excess unreacted dye was removed by two rounds of ethanol precipitation. Solid pellet was redissolved in water (0.1 mM) and stored at −20°C. To assemble the DNA construct, 2 µL of 100 µM Cy3B-labeled non-template strandwith a 28-nt handle was mixed with 3 µL of 100 µM Cy5-labeled template strand and 14 µL of ∼100 mM KGlu (Tris, pH 7.8). The mixture was heated to 95°C for 5 min, then slowly cooled to 22°C at –1°C/min in a PCR thermocycler to promote annealing. Subsequently, 1 µL of 100 µM 28-nt biotin-handle DNA was added and incubated at 30°C for 5 min. Holoenzymes were assembled by incubating 15.5 µL of 11.13 µM core RNAP with 4.5 µL of 116.94 µM σ⁷⁰ at 30°C for 15 min, yielding an approximate 1:3 molar ratio of core RNAP to σ⁷⁰. The resulting holoenzyme (8.625 µM) was diluted to 3.45 µM in storage buffer (20 mM Tris-HCl (pH 7.8), 50 mM NaCl, 3 mM DTT), aliquoted at 1 µL, and stored at –80°C. DNA constructs were diluted to 10 pM in washing buffer (100 mM potassium glutamate, 10 mM MgCl₂, 0.2 mg/mL BSA, 1 mM cysteine HCl, 1 mM DTT, 20 mM Tris-HCl (pH 7.8)) and introduced into a NeutrAvidin-pretreated flow chamber prepared from PEG/biotin-PEG–coated quartz slides and coverslips. After 2 min, excess sample was removed by washing buffer, leaving DNA constructs immobilized via a biotin–NeutrAvidin–biotin linkage. RNAP titrations were carried out in imaging buffer (3.46 mM protocatechuic acid (PCA), 0.315 U/mL protocatechuate-3,4-dioxygenase (PCD), 1.6 mM Trolox, 100 mM potassium glutamate, 10 mM MgCl₂, 0.2 mg/mL BSA, 1 mM cysteine HCl, 1 mM DTT, 20 mM Tris-HCl (pH 7.8), and 0–15 nM RNAP). Data acquisition was performed on a home-built total internal reflection fluorescence (TIRF) microscope as described previously ^94^. For each condition, 30 short movies (∼30 frames at 30 fps) were recorded, and averaged images (10 frames) were generated for FRET analysis ^95^. Single-molecule spots were identified, and Cy3B/Cy5 fluorescence intensities were integrated to calculate FRET efficiencies, which were compiled into histograms. Additionally, long movies (∼1000 frames at 10 fps) were collected to monitor dynamics. Single-molecule trajectories were analyzed in the same way, with Cy3B and Cy5 intensities and corresponding FRET values recorded over time.

## Data Availability

The raw sequence reads of promoter variant libraries are deposited under National Center for Biotechnology Information BioProject ID PRJNA1049603. The codes of the promoter prediction models are available at Zenodo (https://doi.org/10.5281/zenodo.17577923) and GitHub (https://github.com/AntonySTKuo/Biophysical_Promoter_Model).

## Supplementary Information

Supplementary Information (2 items) is available online:

**Supplementary Materials**, including Supplementary Fig. S1-S8, Supplementary Table S1-S6, Supplementary Data S1 caption, and Supplementary References

**Supplementary Data S1.** Promoter sequences and strength of PL_TetR_, PL_LuxR_, and PL_CueR_ variants. The Excel file contains three tables, each listing the −35 and −10 sequences and the basal and induced expression levels of promoter variants, quantified from two sort-seq replicates.

## Author Contributions

Conceptualization: H.H.D.C.; Methodology: I.R.L., S.T.K.; Investigation: S.T.K., W.Y.S., S.W.L., C.W.N., C.C.C., Z.P., H.H.D.C.; Formal analysis: S.T.K., I.R.L., Z.P., C.W.N.; Visualization: S.T.K., C.C.C., I.R.L., H.H.D.C.; Supervision: N.L.C., L.W.Y., I.R.L., H.H.D.C.; Funding acquisition: N.L.C., L.W.Y., I.R.L., H.H.D.C.; Writing – Original Draft: S.T.K., N.L.C., I.R.L., H.H.D.C.; Writing – Review & Editing: N.L.C., Z.P., L.W.Y., I.R.L., H.H.D.C.

## Acknowledgements

We thank the National Taiwan University College of Life Science Technology Commons for instrument support, National Center for High-performance Computing and National Institutes of Applied Research for providing computational and storage resources, and Ben-Yang Liao, Hsuan-Chen Wu, and Chao-Ping Hsu for insightful discussions.

